# Predicting plant biomass accumulation from image-derived parameters: Image-derived parameters predicting plant biomass

**DOI:** 10.1101/046656

**Authors:** Dijun Chen, Rongli Shi, Jean-Michel Pape, Christian Klukas

## Abstract

Image-based high-throughput phenotyping technologies have been rapidly developed in plant science recently and they provide a great potential to gain more valuable information than traditionally destructive methods. Predicting plant biomass is regarded as a key purpose for plant breeders and ecologist. However, it is a great challenge to find a suitable model to predict plant biomass in the context of high-throughput phenotyping. In the present study, we constructed several models to examine the quantitative relationship between image-based features and plant biomass accumulation. Our methodology has been applied to three consecutive barley experiments with control and stress treatments. The results proved that plant biomass can be accurately predicted from image-based parameters using a random forest model. The high prediction accuracy based on this model, in particular the cross-experiment performance, is promising to relieve the phenotyping bottleneck in biomass measurement in breeding applications. The relative contribution of individual features for predicting biomass was further quantified, revealing new insights into the phenotypic determinants of plant biomass outcome. What’s more, the methods could also be used to determine the most important image-based features related to plant biomass accumulation, which would be promising for subsequent genetic mapping to uncover the genetic basis of biomass.

**One-sentence Summary:** We demonstrated that plant biomass can be accurately predicted from image-based parameters in the context of high-throughput phenotyping.

**Footnotes:** This work was supported by the Leibniz Institute of Plant Genetics and Crop Plant Research (IPK), the Robert Bosch Stiftung (32.5.8003.0116.0) and the Federal Agency for Agriculture and Food (BEL, 15/12-13, 530-06.01-BiKo CHN) and the Federal Ministry of Education and Research (BMBF, 0315958A and 031A053B). This research was furthermore enabled with support of the European Plant Phenotyping Network (EPPN, grant agreement no. 284443) funded by the FP7 Research Infrastructures Programme of the European Union.

## Introduction

Biomass accumulation is an import indicator of crop final product and plant performance. It is thus considered as a key trait in plant breeding, agriculture improvement and ecological applications. The conventional approach of measuring plant biomass is very time consuming and labour intensive since plants need to be harvested destructively to obtain the fresh or dry weight. Moreover, the destructive method makes multiple measurements of the same plant over time impossible. With the development of new technology, digital image analysis has been used more broadly in many fields, as well as in plant research. It allows faster and more accurate plant phenotyping and has been proposed as an alternative way to infer plant biomass.

In recent years, plant biomass has been subject to intensive investigation by using high-throughput phenotyping (HTP) approaches in both controlled growth chambers (Tackenberg, 2007, Golzarian et al., 2011, Feng et al., 2013) and field environments (Ehlert et al., 2008, Ehlert et al., 2010, Erdle et al., 2011, Busemeyer et al., 2013, Cao et al., 2013), demonstrating that the ability of imaging-based methods to infer plant biomass accumulation. On the other hand, to produce reliable assessments, suitable model types needs to be established and model construction requires integration of many components such as efficient mathematical analysis and representative data. While there are some developed models for predicting plant biomass, most of them have certain limitations. For example, Golzarian *et al*. (2011) modelled the plant biomass (dry weight) in wheat (*Triticum aestivum* L.) as a linear function of projected area, assuming plant density was constant. However, this method under-estimated dry weight of salt stressed plants and overestimated that of control plants. Although the authors argued that the bias was largely related to plant age and the model might be improved by including the factor of plant age (Golzarian et al., 2011), the differences in plant density between stressed and control plants may be caused by different physiological properties of plants rather than plant age. In another study, Busemeyer *et al*. (2013) developed a calibrated biomass determination model for triticale (x *Triticosecale* Wittmack L.) under field conditions based on multiple linear regression analysis of a diverse set of parameters, considering both, the volume of the plants and their density. Indeed, this model largely improved the prediction accuracy of the calibration models based on a single type of parameters and can precisely predict biomass accumulation across environments (Busemeyer et al., 2013). Another concern is that the number of traits used in these studies were quite limited and perhaps not representative enough. Therefore, a more effective and powerful model is needed to overcome these limitations and to allow better utilization of the image-based plant features which are obtained from non-invasive phenotyping approaches.

In this paper we present a general framework for investigating the relationships between plant biomass (referred to as shoot biomass hereafter) and image-derived parameters. We applied a multitude of supervised and unsupervised statistical methods to investigate different aspects of biomass determinants by a list of representative phenotypic traits in three consecutive experiments in barley. The results showed that image-based features can accurately predict plant biomass output and collectively reflect large proportions of the variation in biomass accumulation. We elucidated the relative importance of different feature categories and of individual features in prediction of biomass accumulation. The differences in the contribution of the image-based features for prediction of two types of biomass measurements, fresh weight and dry weight were compared as well. Furthermore, our models were tested for the possibility of predicting plant biomass in different experiments with different treatments. As high-throughput plant phenotyping is a technique which is becoming more and more widely used for automated phenotype in plant research, especially in plant breeding, we anticipate that the methodologies proposed in this work will have various potential applications.

## Results

### Development of statistical models for modelling plant biomass accumulation using image-based features

In the previous studies (Klukas et al., 2014, Chen et al., 2014), we have shown that a single phenotypic trait - the three-dimensional digital volume, which is a derived feature from projected side and top areas - can be reasonably predictive to estimate plant biomass accumulation. We expect that the predictive power could be improved when multiple phenotypic traits are combined in a prediction model since plant biomass is determined not only by their structural features but also by their density (physiological properties). To further investigate the relationship between image-derived parameters and plant biomass accumulation, deep phenotyping data which contain both structural (e.g., geometric traits) and physiological traits (e.g., plant moisture content; **Fig. 1,A** and **B**) were analysed.

**Figure 1.**
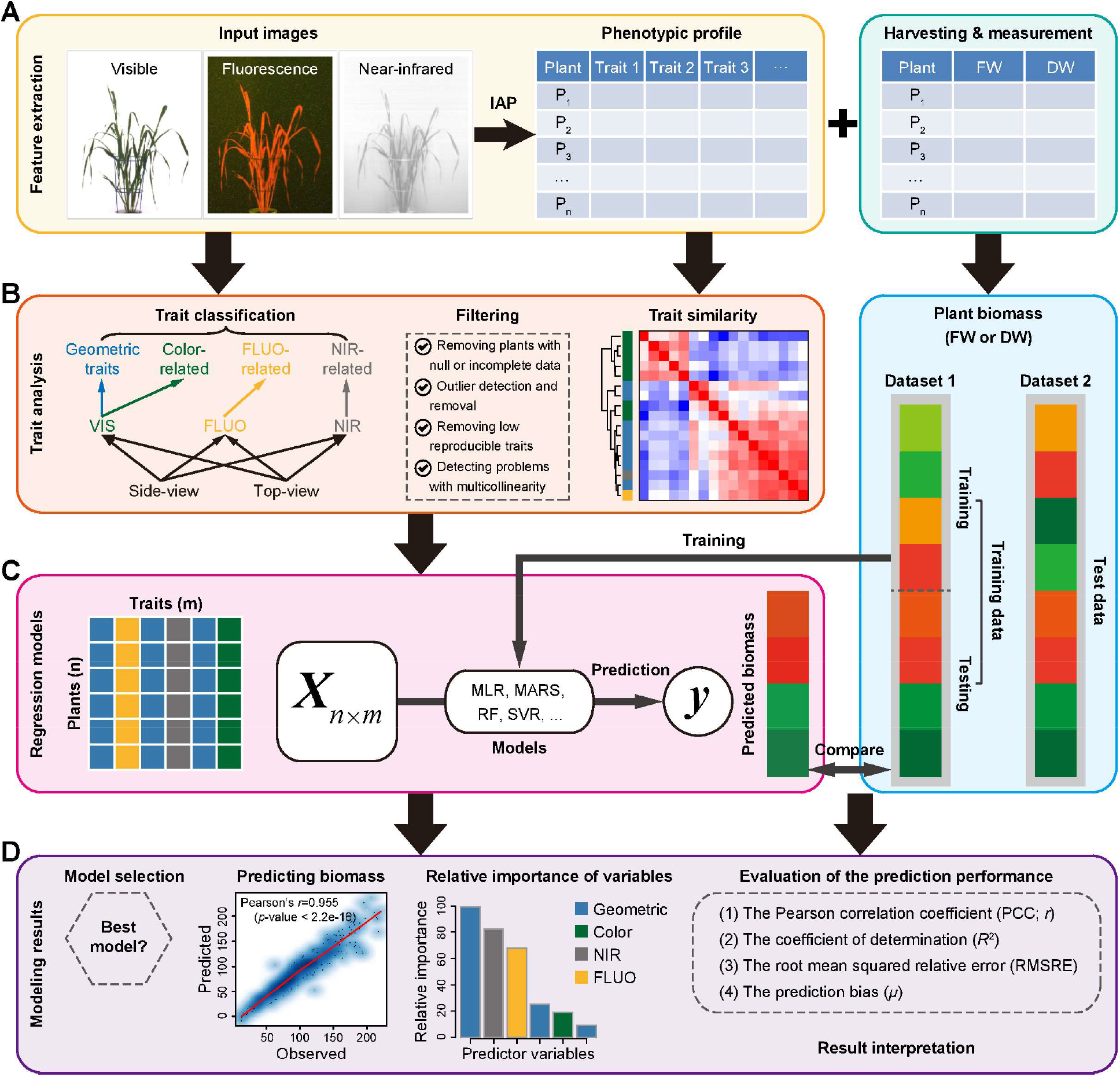
**Modeling pipeline for predicting plant biomass accumulation based on image-derived parameters**. (**A**) Input data, including high-throughput image data and manually measured biomass data. Plants were phenotyped using various cameras such as visible (or color), fluorescence (FLUO) and near-infrared (NIR) sensors. Image analysis was performed with IAP software for feature extraction. The same plants were harvested and measured at the end of growth stage. Generally, two types of biomass was measured: fresh weight (FW) and dry weight (DW). (**B**) Trait processing. All the phenotypic traits are grouped into four categories: geometric, color-related, FLUO-related and NIR-related traits. Phenotypic data were subjected to quality check to remove low-quality data. (**C**) Each plant was described by a list of traits, resulting in a predictor matrix whose rows represent plants and columns represent image-based traits. This matrix was used to predicted plant biomass accumulation by MLR (multivariate linear regression), MARS (multivariate adaptive regression splines), RF (random forest) and SVR (support vector regression) models. The right panel represents the schema of model validation. In the first schema, a dataset (Dataset 1) was divided into training set and testing set in a ten-fold cross-validation manner. In the second schema, the whole of one dataset (Dataset 1) was used for training and another dataset (Dataset 2) was used for testing. (**D**) Model selection, evaluation and result interpretation. The correlation of the predicted values and measured values was used to assess the overall performance of the model.

Models were constructed to quantify the ability of imaging-based features to statistically predict the biomass accumulation. The models were developed by using four widely used machine-learning methods (**Fig. 1C**): multivariate linear regression (MLR), multivariate adaptive regression splines (MARS), random forest (RF) and support vector regression (SVR), which have extensively been used in accurate prediction of gene expression (Cheng et al., 2012, Cheng & Gerstein, 2012, Cheng et al., 2011, Dong et al., 2012, Karlić et al., 2010) and DNA methylation levels (Ma et al., 2014, Zhang et al., 2015, Das et al., 2006, Zheng et al., 2013). We combined the biomass measurements (fresh weight [FW] and/or dry weight [DW]) with image-based features and then divided them into a training data set and a test data set. A model was trained on the training data set and has then been applied to the test data set to predict the plant biomass. The relationship between plant biomass accumulation and image-based features was assessed based on the criterion of the Pearson correlation coefficient (*r*) between the predicted values and the actual values, or the coefficient of determination (*R^2^*; the percentage of variance of biomass explained by the model; **Fig. 1D**).

Our methodology was applied to three consecutive experiments (**Fig. 2A**; **Supplemental Table S1** and **Data S1**), which were designed to investigate vegetative biomass accumulation in response to two different watering regimes under semi-controlled greenhouse conditions in a core set of barley cultivars by non-invasive phenotyping (Chen et al., 2014, Neumann et al., 2015). There were 312 plants with 18 genotypes for each experiment. Plants were monitored using three types of sensors (visible, fluorescence [FLUO] and near-infrared [NIR]) in a LemnaTec-Scanalyzer 3D imaging system. An extensive list of phenotypic traits ranging from geometric (shape descriptors) to physiological properties (i.e., colour-, FLUO- and NIR-related traits) could be extracted from the image data (**Fig. 1B**) using our image processing pipeline IAP (Klukas et al., 2014). A representative list of traits for each plant in the last growth day were selected to test their ability to predict plant biomass.

**Figure 2.**
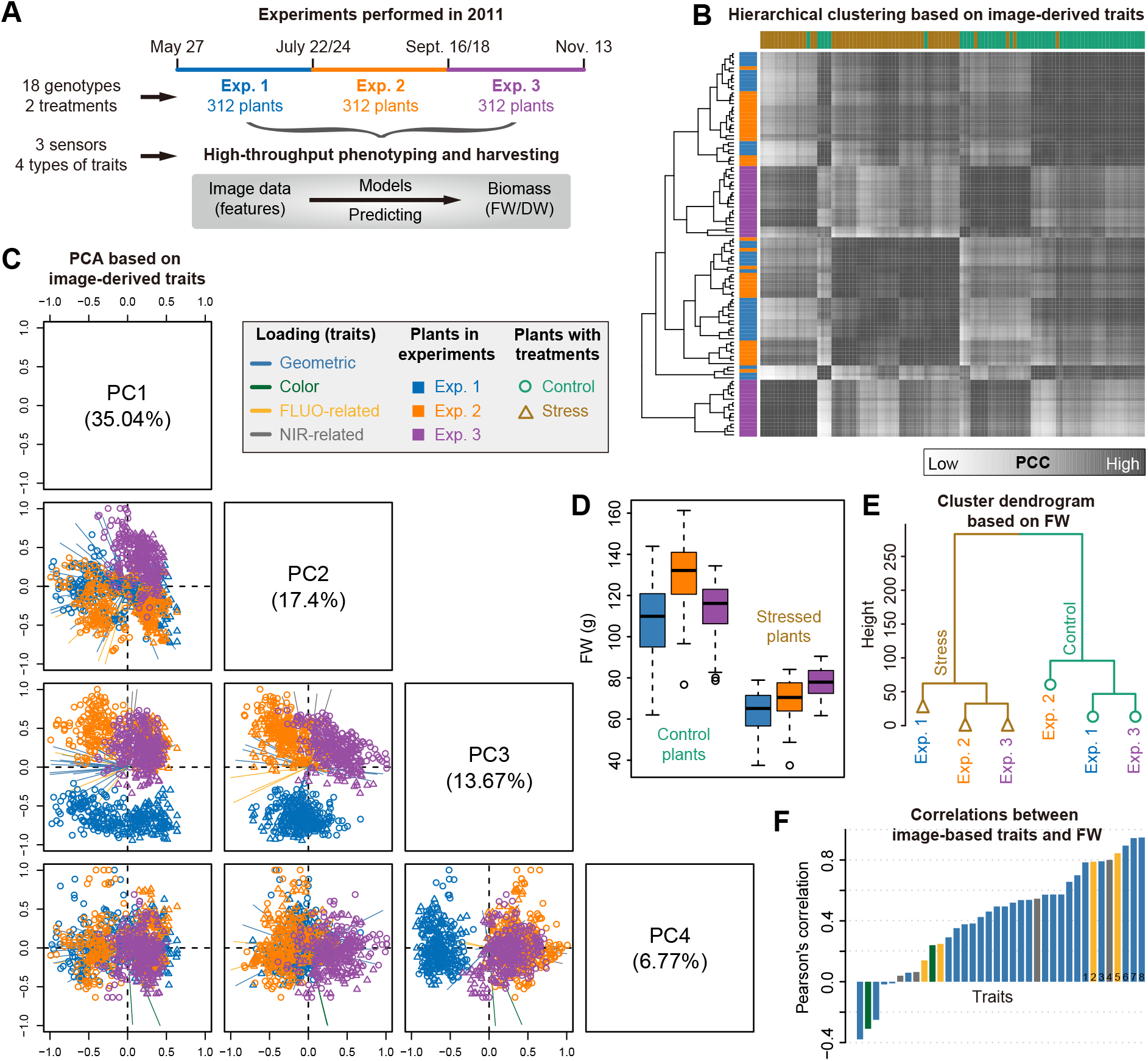
**Predictability of image-based traits to plant biomass**. (**A**) Schema depicting three consecutive high-throughput phenotyping experiments in barley. Plants in each experiment were harvested for biomass measurements: fresh weight (FW; for all experiments) and dry weight (DW; only for experiment 1). (**B**) Heatmap of Pearson’s correlations between plants. Pearson’s correlation coefficient (PCC) was calculated based on image-derived traits. Cluster dendrograms for experiments (left) and treatments (top) are shown. (**C**) Scatter plots showing projections of top four Principal components (PCs) based on PCA of image-based data. The component scores (shown in points) are colored and shaped according to the experiments (as legend listed in the box). The component loading vectors (represented in lines) of each traits (as colored according to their categories) were superimposed proportionally to their contribution. (**D**) Boxplot showing the distribution of FW across different experiments. (**E**) A dendrogram from cluster analysis based on the means of FW data over genotypes. (**F**) Pearson’s correlation (mean values in the three datasets) between image-based traits and FW. Traits with the largest mean correlations values are labeled: 1 — sum of leaf length (side view), 2 — sum of FLUO intensity (side), 3 — plant area border length (side), 4 — sum of NIR intensity (top), 5 — sum of FLUO intensity (top), 6 — projected area (top), 7 — projected area (side) and 8 — digital volume.

### Coordinated patterns of plant-image-based profiles and their relation to plant biomass

We extracted a list of representative and non-redundant phenotypic traits for each plant from image datasets for each experiment (see **Materials and Methods**; **Fig. 1B**). In common for these experiments, overall thirty-six high-quality traits which describe plant growth status in the last growth day were obtained. As a result, each dataset was assigned a matrix whose elements were the signals of different features in different plants (**Fig. 1C**). Unsupervised methods, such as hierarchical clustering (HCA; **Fig. 2B**) and principal component analysis (PCA; **Fig. 2C**) were applied to these datasets. We found that plants from different experiments with different treatments showed clearly distinct patterns of phenotypic profiles. For instance, stressed plants and control plants were separated using PCA by their first principal component (PC1) and also by the top clusters obtained in HCA, while plants from different experiments were distinguished by PC2 and PC3 in PCA or subordinate clusters in HCA. Accordingly, it could be observed that biomass (e.g., FW) of plants from different experiments with different treatments was significantly different (two-way ANOVA, p-value < 2e-16; Fig. 2D). The relationship was reflected by a dendrogram from cluster analysis based on the means of FW over genotypes (Fig. 2E). Furthermore, the overall phenotypic patterns of these plants were similar to their biomass output (Fig. 2, B-E), revealing that these image-based features were potential factors reflecting the accumulation of plant biomass. We thus explored the relationship between the signals of these image-based features and the level of plant biomass output. We calculated the correlation coefficients for each dataset. The correlation patterns were consistent for different datasets and more than half of the features revealed high correlation coefficients (r > 0.5; **Fig. 2F**). Interestingly, both structural features (such as digital volume, projected area and the length of the projected plant area border) and density-related features (such as NIR and FLUO intensities) were involved in the top ranked features.

### Relating image-based signals to plant biomass output

The above analyses suggest that plant biomass can at least be partially inferred from image-based features. To examine which model has the best performance and to select an appropriate model for biomass prediction, we then applied our regression models (**Fig. 1C**) to predict plant biomass using image-based features. Our analyses were focused on the first experiment (i.e., experiment 1), since the phenotypic traits of the corresponding dataset have been intensively investigated in our previous study (Chen et al., 2014). In this experiment, plant biomass was quantified in two forms: FW and DW. We selected a collection of 45 image-derived parameters from this dataset that were non-redundant and highly representative.

We next tried to predict FW (**Fig. 3A**) and DW (**Fig. 3C**) based on this set of image-derived features using four different regression models. The models were respectively tested on control plants, stressed plants and the whole set of plants. The performance of these models was compared and evaluated. Although the performance of these models was roughly similar, RF, SVR and MARS methods had better performance than the MLR method for prediction of both FW (**Fig. 3B**) and DW (**Fig. 3D**), implying a nonlinear relationship between image-based phenotypic profiles and biomass output. The RF model largely outperformed other models especially in predicting biomass of control plants, accounting for the most variance (*R^2^* = 0.85 for FW and *R^2^ =* 0.62 for DW; **Fig. 3, B and D**, left panels) and showed the best prediction accuracy (Pearson’s correlation *r =* 0.93 for FW and *r =* 0.80 for DW; **Fig. 3, B** and **D**, middle panels). The prediction accuracy of our models (the correlation coefficients between the predicted biomass and the actual biomass) was as well compared with the ability of individual features to predict biomass (here, the “digital volume”; Fig. 3, B and D, middle panels). It was found that our models generally showed better prediction power than the single digital volume-based prediction, indicating that additional features improved the predictive power. In this study, we focused on the results from the RF method in the rest of analysis, although results from different methods were highly consistent and led to the same conclusions.

**Figure 3.**
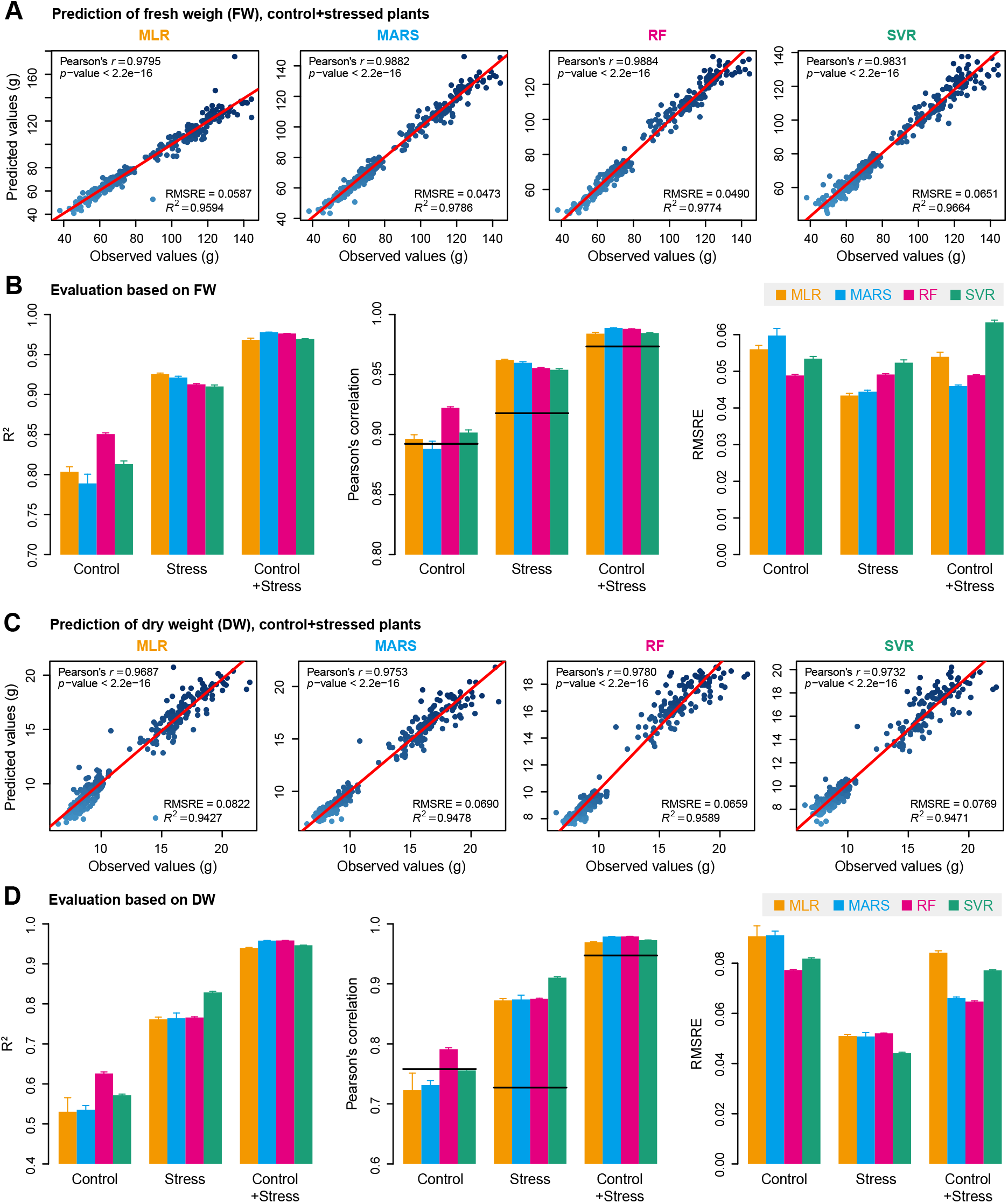
**Quantitative relationship between image-based features and plant biomass**. (**A**) and (**C**) Scatter plots of manually measured plant biomass (fresh weight [FW] and dry weight [DW]) versus predicted biomass values using four prediction models: multivariate linear regression (MLR), mul-tivariate adaptive regression splines (MARS), random forest (RF) and support vector regression (SVR). The red line indicates the expected prediction (*y = x*). The quantitative relationship between image-based features and biomass is evaluated by Pearson’s correlation coefficient (PCC *r* and its corresponding *p*-value), *RMSRE* (root mean squared relative error) and the percentage of variance explained by the models (the coefficient of determination *R*^2^). (**B**) and (**D**) Summary of the predictive power of each regression models. The results are based on ten-fold cross-validation with ten trials. Models were evaluated based on control plants, stressed plants and the whole set of plants. The solid lines in the middle panel represent PCC between digital volume and biomass for specific datasets.

### Relative importance of different image-based features for predicting plant biomass

As mentioned above, the image-based features could be classified broadly into four categories: plant structure properties, colour-related features, NIR signals, and FLUO-based traits (**Fig. 1B**). The last three types of features reflect plant physiological properties and can be considered as plant density-related traits and are thus related to their fresh or dry matter content. For each individual feature or each type of features, we constructed a degenerate model for biomass prediction using the corresponding feature(s) as the predictor (s). We compared the capability of each individual or type of feature for predicting biomass accumulation in the first experiment (i.e., experiment 1). Geometric features showed the most predictive power among the four categories for prediction of both FW and DW, but were slightly less predictive than all features in a full model (**Fig. 4, A and B**). Strikingly, the predictability of other types of features (such as colour-re la ted and FLUO-based traits) was substantial, indicating that these traits may act as unforeseen factors in biomass prediction. In addition, the NIR-based features showed higher predictive capability for FW than for DW in control and stressed plants, revealing NIR signals were import factors in determining FW accumulation.

**Figure 4.**
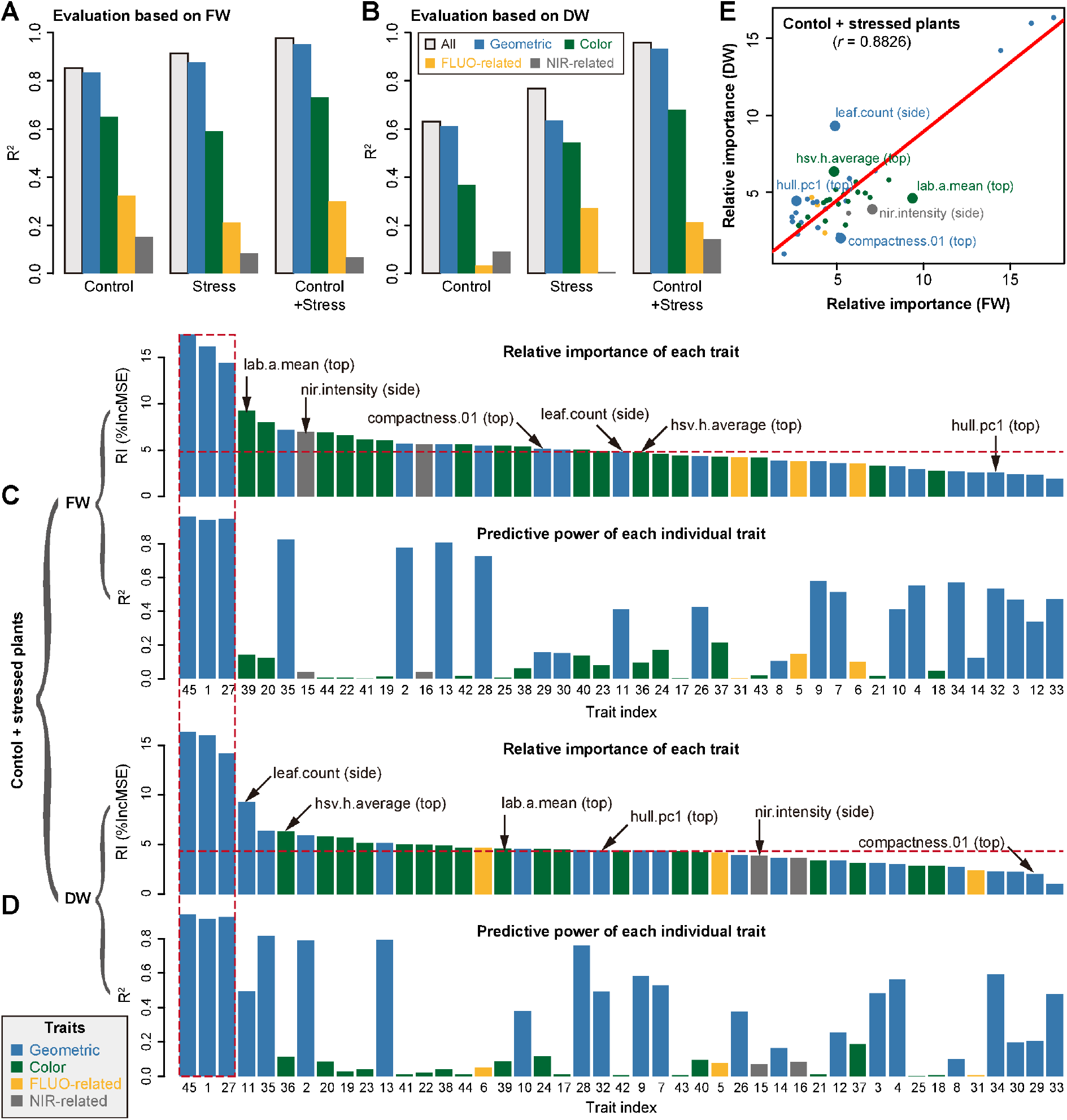
**The relative importance of image-based features in prediction of plant biomass**. The capabilities of different types of image-based features to predict plant biomass based on evaluation of either fresh weight (FW) (**A**) or dry weight (DW) (**B**). The overall predictive accuracies of each types of features are indicated. Grey bar denotes the predictive accuracy using all features. The relative importance of each feature in the Random Forest model (upper panel) and the predictive accuracy of each individual feature as the single predictor (lower panel) based on investigation of either FW (**C**) or DW (**D**). The calculation is based on the whole set of plants (control and stressed plants). Note that feature labels are shared in the upper and lower panels. Features are shown in numbers as ordering by their names. Three features highlighted in red dash box are digital volume, projected side area and projected top area. (**E**) Comparison of the relative importance of features in prediction of FW and DW. The top six most different features are highlighted and labeled.

Next, we investigated the relative importance (RI) of each feature for predicting biomass using a full model in the whole set of plants (i.e., “control + stressed plants”; **Fig. 4, C** and **D**, upper panels). In a RF model, the RI of a feature is calculated as the increase of prediction error (%IncMSE) when phenotypic data for this feature is permuted (Breiman, 2001), and thus indicates the contribution of the feature after considering its intercorrelation in a model. We found that the top ten most important features in the full model for predicting FW and DW included both structure and density-related traits. As expected, projected area (from side or top view) and digital volume were the top ranked features, which have individually been considered as proxies of shoot biomass in previous studies (Dietz & Steinlein, 1996, Leister et al., 1999, Paruelo et al., 2000, Walter et al., 2007, Arvidsson et al., 2011, Golzarian et al., 2011, Chen et al., 2014, Hairmansis et al., 2014, Neilson et al., 2015).

In principle, we would expect that highly important features in the full model would be related to a high predictive power in a degenerate model. Surprisingly, there was no clear correlation observed between the feature importance and their predictive power (Fig. 4, C and D). For example, several colour-related and NIR-based features which were in the top ten list of the most important features revealed insubstantial predictive power in individual models. This observation implies that the relation of the underlying biomass determinants is extremely complex and not a linear combinations of the investigated features.

Furthermore, we compared the relative importance of each feature in predicting FW and DW (**Fig. 4E**). Although a positive correlation (*r =* 0.88) between the feature importance for FW and DW could be observed, several features showed large differences in their ability to interpret FW or DW, including “nir.intensity” (derived from side view images), “compactness.01” (top), “hull.pc1” (top), “leaf.count” (side), “hsv.h.average” (top) and “lab.a.mean” (top). For instance, NIR intensity and plant compactness (top view) may be important for predicting FW but not for DW. We also performed the above analyses by using only control (**Supplemental Fig. S1**) or stressed plants (**Supplemental Fig. S2**), respectively. We found that the patterns of feature importance were distinct between these two groups of plants. For example, NIR intensity was ranked as the top fifth feature for predicting FW for stressed plants but was not substantially important for control plants. These findings suggest that there are differences in underlying plant biomass determinants in these kinds of treatment situations that are also reflected by their image-based phenotypic traits.

### Image-based features are predictive of plant biomass across experiments with similar conditions or treatments

In order to explore whether our models were generalizable across different experiments, we applied our models trained in one experiment to predict biomass (herein FW) in other experiments using a common set of features. Examples of such cross-experiment predictions are shown in **Figure 5A**. We tested and illustrated all possibilities for cross prediction using the whole set of plants in the corresponding experiment. In general, the prediction accuracy within individual experiments remained high (*r >* 0.97 and *R^2^ >* 0.93 for all three experiments; **Fig. 5B**), revealing that our models were effectively predicting plant biomass based on image-derived feature signals among different experiments. Moreover, the prediction accuracy for cross-experiment prediction was still relatively high, with *r >* 0.81 and *R^2^ >* 0.65, implying that our models accurately captured the relationships among the various image-based features. However, we observed that the third experiment had relative weaker correlations with the other two experiments for predicting biomass, while the first two experiments showed strong correlations or even nearly identical results when being compared with each other (**Fig. 5A**). This might be related mainly to seasonal (temperature and illumination) differences which caused different plants behaviours in experiment 3 as explained by the authors (Neumann et al., 2015).

**Figure 5.**
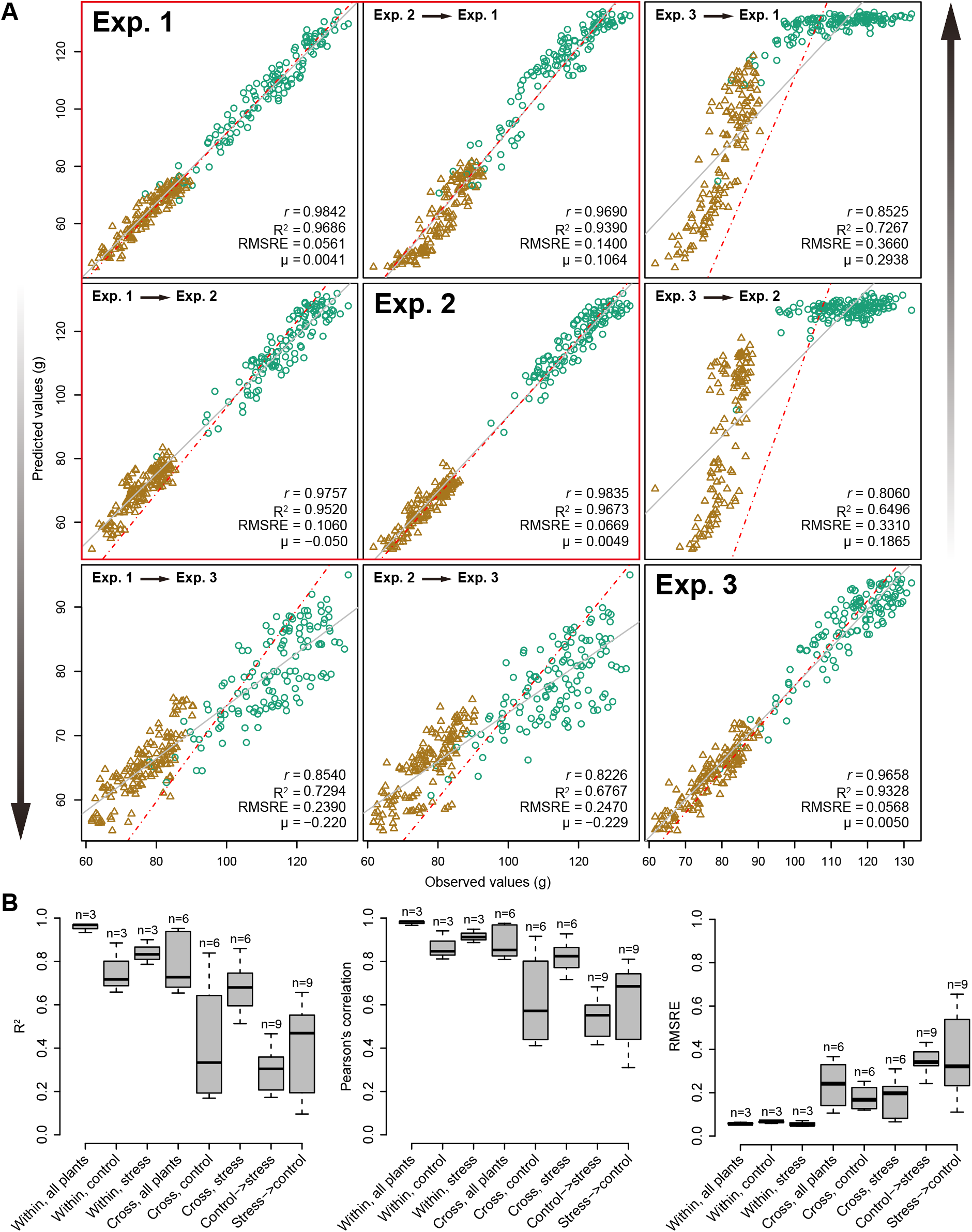
**Comparison of prediction accuracy across different experiments**. (**A**) Application of the model learned from one experiment to other experiments. (**B**) Boxplots of coefficient determination (*R*^2^, left) Pearson’s correlation coefficients (*r*, middle) and the root mean squared relative error (*RMSRE*, right) for different comparisons. “Within” denotes a model trained and tested on data from the same dataset with specific treatments (control, stress or both), and “Cross” represents a model trained on one dataset and tested on another dataset. “Control *→* stress” denotes a model trained on data with control treatment and tested on data with stress treatment, and vice versa for “stress *→* control”.

At the same time, we tested cross predictability of our models using treatment-specific data in the experiments (**Fig. 6**). Similar results were obtained as above using the whole dataset (**Fig. 5B**). The weak predictive power for cross-prediction involving control plants from the third experiment was most clearly observable in the low accuracy in the biomass prediction of this particular subset of plants. Generally, control and stressed plants were found to have very weak predictive power when related to each other (**Fig. 6**), as also supported by the distinct patterns of relative feature importance between these two plant groups (**Supplemental Figs. S1** and **S2**).

**Figure 6.**
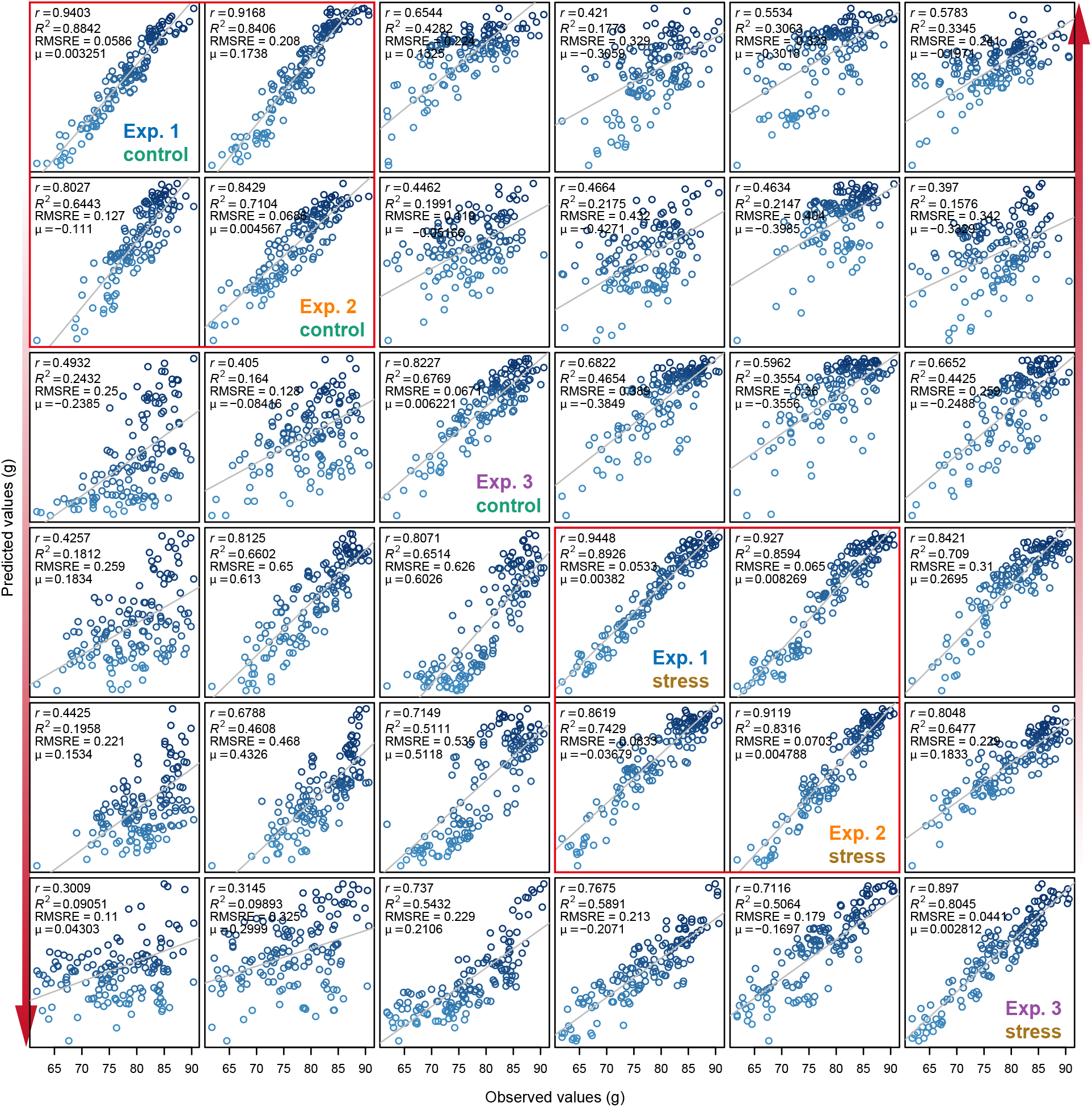
**Comparison of prediction accuracy across different treatments**. Refer to Figure 5B for legend.

## Discussion

Biomass is a complex but important trait in functional ecology and agronomy for studying plant growth, crop productive potential and plant regeneration capabilities. Many different techniques, either destructive or non-destructive, have been used to estimate biomass (Catchpole & Wheeler, 1992). Compared with the traditional destructive methods for measuring biomass, non-destructive imaging methods provide a faster, more accurate approach for plant phenotyping. In recent years, more and more high-throughput plant phenotyping platforms have been set up and applied worldwide. Accordingly, it becomes a current challenge to establish models utilizing the big datasets gained from high-throughput imaging systems. Although previous attempts have been made to estimate plant biomass from image data, most of these studies consider only a single image-based feature or very few features in their models which are often linear-based, ignoring the fact that the phenotypic components underlying biomass accumulation are presumably complex. Accurately predicting biomass from image data requires efficient mathematical models as well as representative image-derived features.

In this study, we have presented a systematic analysis of relationships between plant biomass accumulation and image-derived signals, to confirm the assumption that biomass can be accurately predicted from image-based parameters. We built a random forest model of biomass accumulation using a comprehensive list of representative image-based features. The comparison between the RF model and alternative regression models indicated that the RF model outperforms other models in terms of (1) better predictive power -especially in comparison with the linear model, confirming the complex phenotypic architecture of biomass, and (2) feasible biological interpretability - the ability to readily extract information about the importance of each feature in prediction. The high prediction accuracy based on this model, in particular the cross-experiment performance, is promising to relieve the phenotyping bottleneck in biomass measurement in breeding applications. For example, based on an established small reference dataset which is used to train a RF model, it is possible to predict biomass in several large plant populations within one experiment or across several experiments using image data by taking advantage of high-throughput phenotyping technologies. However, considering the environmental effects on biomass accumulation, the application of our model will require the testing experiments showing similar conducted conditions with that of the reference experiments. Alternatively, the model can be trained from a much larger reference panel of plants that are grown in diverse environmental conditions which is then is applied to a diverse set of experiments. The first evidence for this notion is the observation that our model showed more predictive power in plants with two treatments than with a single treatment (**Fig. 3**). Indeed, when applying our model to the combined dataset from all the three experiments, we found the prediction accuracy remains very high (*R^2^* = 0.96 and *r =* 0.98, average values from ten times of ten-fold cross-validation). Another approach to improve applicability of models, which could not be tested in this study, would be to improve the data base for the training, by acquiring data from additional environment sensors. Temperature, humidity, and illumination data would certainly help to explain differences in the growth patterns among experiments, performed in different growth seasons. To this end, we expect that our approach is extensible by incorporating such sensor data in the data matrices.

In contrast to previous studies (Dietz & Steinlein, 1996, Leister et al., 1999, Paruelo et al., 2000, Walter et al., 2007, Arvidsson et al., 2011, Golzarian et al., 2011, Hairmansis et al., 2014, Neilson et al., 2015, Tackenberg, 2007, Feng et al., 2013), in which biomass was investigated using only single image-derived parameter (such as projected area) or several geometric parameters, our analyses extended these studies by incorporating more representative features that cover both structural and physiological-related properties into a more sophistic model. Although the predictive power of our model is roughly higher than that of single feature-based prediction, such as the digital volume (**Fig. 3**) (Chen et al., 2014), our model also reveals the relative contribution of individual feature in prediction of biomass. The information regarding the importance of each feature will offer new insights into the phenotypic determinants of plant biomass outcome. Interestingly, we found that several top ranked features, such as digital volume and NIR intensity, showed genetic correlations with biomass of fresh weight (**Fig. 4C**) (Chen et al., 2014), implying these top ranked features may represent the main “phenotypic components” of biomass outcome and can be further used to dissect genetic components underlying biomass accumulation. As image-based high-throughput phenotyping in plants developed mainly in recent years and therefore few corresponding modelling studies have been performed, we believe that our model could be further improved when new types of cameras and/or newly defined features are available.

In summary, we have developed a quantitative model for dissecting the phenotypic components of biomass accumulation based on image data. Apart from predicting biomass outcome, the methods can be used to determine the most important image-based features related to plant biomass accumulation, which are promising for subsequent genetic mapping to uncover the genetic basis of biomass. We anticipate that these statistical methods will be broadly used in plant breeding in the context of phenomics.

## Materials and Methods

### Germplasm and experiments

Barley plant image data were obtained as described previously (Chen et al., 2014, Neumann et al., 2015). Briefly, a core set of 16 two-rowed spring barley cultivars (*Hordeum vulgare* L.) and two parental cultivars of a double haploid (DH) were monitored for vegetative biomass accumulation. Three independent experiments with identical setup were performed in a (semi-) controlled greenhouse at IPK by using the automated phenotyping and imaging platform LemnaTec-Scanalyzer 3D. Experiments were performed consecutively from May to November 2011 over a period of 58 days each (**Supplemental Table S1**). The greenhouse setup enabled sowing for the next experiment already 2 days before the old experiment ended. For this, new pots were placed in the middle of the greenhouse, while the old experiment was still on the conveyer belts.

Each experiment consisted of two treatments: well-watered (control treatment) and water limited (drought stress treatment). In each treatment, nine plants per core set cultivar as well as six plants per DH parent were tested. This resulted in a total of 312 plants per experiment, corresponding to the maximal capacity of the phenotyping platform. Watering and imaging were performed daily. Drought stress was imposed by intercepting water supply from 27 days after sowing (DAS 27) to DAS 44. Stressed plants were re-watered at DAS 45. In total, for each of the experiments about 100 GB of raw (image) data was accumulated. At the end of experiments (DAS 58), plants were harvested to measure above-ground biomass in form of plant fresh weight (FW; for all experiments) and/or dry weight (DW; for experiment 1).

**Image analysis**

Image datasets were processed by the barley analysis pipelines in the IAP software. Analysed results were exported in the csv file format via IAP functionalities, which can be used for further data inspection. The result table includes columns for different phenotypic traits and rows as plants are imaged over time. The corresponding metadata is included in the result table as well.

Each plant was characterized by a set of phenotypic traits also referred to as features, which were grouped into four categories: geometric features, fluorescence-related (FLUO-related) features, colour-related features and near-infrared-related (NIR-related) features. These traits were defined by considering image information from different cameras (visible light, fluorescence and near infrared) and imaging views (side and top views). See the IAP online documentation (http://iapg2p.sourceforge.net/documentation.pdf) for details about trait definition.

### Feature selection

Feature selection was performed with the same procedure as described in (Chen et al., 2014). We applied the feature selection technique to each dataset. Generally, we captured almost identical subset features from different datasets. We manually added several representative traits due to removal by variance inflation factors. For example, the digital volume and projected area are highly correlated with each other but we kept both of them, because we would investigate the predictive power of both features. Moreover, the regression models we used are insensitive to collinear features. We thus kept as much representative features as possible. To apply the prediction models among different datasets, a common set of features supported by all the datasets was used.

### Data transformation

Each plant can be presented by a representative list of phenotypic traits, resulting in a matrix ***X_nxm_*** for each experiment, where **n** is the number of plants and ***m*** is the number of phenotypic traits. Missing values were filled by mean values of other replicated plants. To make the image-derived parameters from diverse sources comparable, we normalized the columns of **A”** by dividing the values with the maximum value of each column across all plants. Plants with empty values of manual measurements (FW and DW) were discarded for analysis. These transformed data sets were subjected to regression models.

### Hierarchical clustering analysis and PC A

Hierarchical clustering analysis (HCA) and principle component analysis (PCA) were performed on the transformed data matrix ***X_nxm_*** in the same way as described in (Chen et al., 2014). We also performed HCA using the genotype-level mean value of FW data to check the similarity of overall plant growth patterns in different experiments.

### Models for predicting plant biomass

To understand the underlying relationship between image-derived parameters and the accumulated biomass (such as FW and DW), we constructed predictive models based on four different machine-learning methods: multivariate linear regression (MLR), multivariate adaptive regression splines (MARS), random forest (RF) and support vector regression (SVR). In these models, the normalized phenotypic profile matrices ***X_nxm_*** for a representative list of phenotypic traits were used as predictors (explanatory variables) and the measured DW/FW as the response variable ***Y***.

All these models were implemented in R (http://www.r-project.org/; release 2.15.2). To assess the relative contribution of each phenotypic trait to predicting the biomass. We also calculated the relative feature importance for each model. Specifically, for the MLR model, we used the “lm” function in the base installation packages. The relative importance of predictor variables in the MLR model was estimated by a heuristic method (Johnson, 2000) which decomposes the proportionate contribution of each predictor variable to *R^2^*. For MARS, we used the “earth” function in the *earth* R package. The “number of subsets (nsubsets)” criterion (counting the number of model subsets that include the variable) was used to calculate the variables feature importance, which is implemented in the “evimp” function. For the RF model, we used the *randomForest* R package which implements Breiman’s random forest algorithm (Breiman, 2001). We chose the “%IncMSE” (increase of mean squared error) to represent the criteria of relative importance measure. For SVR, we utilized the *e1071* R package which provides functionalities to use the *libsvm* library (Chang & Lin, 2011). The absolute values of the coefficients of the normal vector to the “optimal” hyperplane can be considered as the relative importance of each predictor variable contributing to regression (Loo et al., 2007, Iyer-Pascuzzi et al., 2010).

### Evaluation of the prediction models

To evaluate the performance of the predictive models, we adopted a 10-fold cross-validation strategy to check the prediction power of each regression model. Specifically, each dataset was randomly divided into a training set (90% of plants) and a testing set (10% of plants). We trained a model on the training data and then applied it to predict biomass for the testing data. Afterwards, the predicted biomass in the testing set was compared with the manually measured biomass. The predictive accuracy of the model can be measured by

1. the Pearson correlation coefficient (PCC; *r*) between the predicted values and the observed values;
2. the coefficient of determination (*R^2^*) which equals to the fraction of variance of biomass explained by the model, defined as

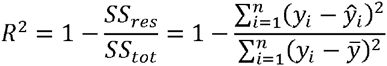

where *SS_res_* and *SS_tot_* are the sum of squares for residuals and the total sum of squares, respectively, *y_L_* the predicted and *y_t_* the observed biomass of the *i*th plant, *y* is the mean value of the observed biomass; and

1. the root mean squared relative error of cross-validation, defined as

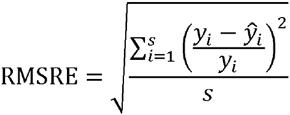

where *s* denotes the sample size of the testing dataset.

We repeated the cross-validation procedure ten times. The mean and standard deviation of the resulting *R^2^* and RMS RE values were calculated across runs.

To illustrate the broad utility of our methods across seasons (thus different growth environments) and treatments (e.g., control versus drought stress) in the same season, we applied the models in different contexts with cohort validation. Specifically, we trained the biomass prediction models under one specific context and predicted biomass in another different context and *vice versa*. The predictive accuracy of the model was evaluated based on the measures *R^2^* and RMSRE as described above. Furthermore, the predictive power was reflected by the bias *μ* between the predicted and observed values, defined as

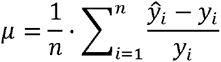

where *n* denotes the sample size of the dataset. This bias indicates over- (μ > 0) or underestimation (μ < 0) of biomass.

## Acknowledgements

We would like to thank Ingo Mücke for his management of the LemnaTec system operations. We thank Dr. Kerstin Neumann for providing the valuable plant data. We thank Michael Ulrich for performing software tests and helping in data analysis.

## Author contributions

D.J.C. designed the research, C.K. supervised the project. J.M.P. and C.K. analyzed image data. D.J.C. analyzed the image analysis result data and wrote the article. R.L.S edited the article and contributed partly for writing. All authors read and approved the final version of the article.

## Supplemental Data

The following supplemental materials are available.

**Supplemental Figure S1**. The relative importance of image-based features in prediction of biomass in control plants. Refer to **Figure 4** for legend. The calculation was based on control plants.

**Supplemental Figure S2**. The relative importance of image-based features in prediction of biomass in stressed plants. Refer to **Figure 4** for legend. The calculation was based on stressed plants.

**Table S1.** Overview of three high-throughput phenotyping experiments in barley.

| Experiment | #plants/#genotypes<sup>1</sup> | Date of sowing | Date of harvesting | Biomass<sup>2</sup> |
| --- | --- | --- | --- | --- |
| Exp. 1 | 310/18 | 27.05.2011 | 24.07.2011 | FW & DW |
| Exp. 3 | 310/18 | 22.07.2011 | 18.09.2011 | FW |
| Exp. 3 | 309/18 | 16.09.2011 | 13.11.2011 | FW & DW |

<sup>1</sup>Number of plants or genotypes used in analysis (filtered data).
<sup>2</sup>Types of biomass measurement. FW: fresh weight; DW: dry weight.

**Supplemental Data S1**. Manual data and image-derived data in the three experiments.

